# Computational detection, characterization, and clustering of microglial cells in a mouse model of fat-induced postprandial hypothalamic inflammation

**DOI:** 10.1101/2025.02.04.636433

**Authors:** Clara Sanchez, Morgane Nadal, Céline Cansell, Sarah Laroui, Xavier Descombes, Carole Rovère, Éric Debreuve

## Abstract

Obesity is associated with brain inflammation, glial reactivity, and immune cells infiltration. Studies in rodents have shown that glial reactivity occurs within 24 hour of high-fat diet (HFD) consumption, long before obesity development, and takes place mainly in the hypothalamus (HT), a crucial brain structure for controlling body weight. Understanding more precisely the kinetics of glial activation of two major brain cells (astrocytes and microglia) and their impact on eating behavior could prevent obesity and offer new prospects for therapeutic treatments. To understand the mechanisms pertaining to obesity-related neuroinflammation, we developed a fully automated algorithm, NutriMorph. Although some algorithms were developed in the past decade to detect and segment cells, they are highly specific, not fully automatic, and do not provide the desired morphological analysis. Our algorithm cope with these issues and performs the analysis of cells images (here, microglia of the hypothalamic arcuate nucleus), and the morphological clustering of these cells through statistical analysis and machine learning. Using the k-Means algorithm, it clusters the microglia of the control condition (healthy mice) and the different states of neuroinflammation induced by high-fat diets (obese mice) into subpopulations. This paper is an extension and re-analysis of a first published paper showing that microglial reactivity can already be seen after few hours of high-fat diet (Cansell et al., 2021). Thanks to NutriMorph algorithm, we unravel the presence of different hypothalamic microglial subpopulations (based on morphology) subject to proportion changes in response to already few hours of high-fat diet in mice.

## 1. Introduction

Overweight and obesity are major public health issues respectively affecting more than 39% and 13% of humanity, including 40 million of children (WHO, 2016). Excess bodyweight is associated with a large range of comorbidities, such as cancer and cardiovascular diseases (Hotamisligil, 2006), making it the world fifth leading risk factor for death, with over 4 million fatalities a year (WHO). Studies on animal models and humans show that high-fat diets (HFD) trigger chronic inflammation which affects tissues such as the liver, adipose tissues, skeletal muscles, pancreatic islets, and eventually the central nervous system (CNS) (Lumeng and Saltiel, 2011; Rhea et al., 2017).

The arcuate nucleus (ARC) of the hypothalamus, a regulatory brain region, show significant markers of inflammation (De Souza et al., 2005; Thaler et al., 2012). HFD- induced hypothalamic inflammation has been recently found to deregulate energy homeostasis (Gao and Horvath, 2008; Seong et al., 2019) and to activate glial cells that accentuate inflammatory reactions, leading to reduced satiety (André et al., 2017; Guillemot-Legris et al., 2016; Valdearcos et al., 2017).

It comes across that alterations triggered by diet-induced obesity (DIO) are dependent of the HFD exposition time (Guillemot-Legris and Muccioli, 2017) and involve neuroglial plasticity (Cansell et al., 2021; Nuzzaci et al., 2020). As such, microglia are central players of obesity-induced neuroinflammation. They are extremely sensitive to small changes in their environment, allowing for an hourly response, and are prone to quick morphological changes (Allen and Lyons, 2018; Zhang et al., 2010).

Recent findings suggest that hypothalamic inflammation occurs before the development of obesity. In particular, microgliosis has been found in the ARC after a 72-hour exposure to HFD (Thaler et al., 2012). We have previously shown that neuroinflammation does not only arise due to regular HFD exposure and with the excess bodyweight onset, but is already set on after a few hours of HFD feeding (Cansell et al., 2021). If HFD has deleterious effects on such short time, its regular intake during youth could dramatically foster obesity and its comorbidities in growing adults.

Establishing the kinetic of early activation of microglia in the ARC could offer new prospects for therapeutic treatments. To understand the mechanisms pertaining to obesity-related neuroinflammatory microglia response, we developed a fully automated algorithm: *NutriMorph*. Although some algorithms were developed in the past decade to detect and segment neural cells, they are highly specific, not fully automatic, and do not provide the desired morphological analysis (Liu et al., 2018; Ly et al., 2020; Peng et al., 2010; SheikhBahaei et al., 2018). Our algorithm cope with these issues and performs the analysis of microglial cells images, and the morphological clustering of these cells through statistical analysis and machine learning.

In this paper, we thus detail the *NutriMorph* methodology, which involves microscopy images processing and analysis to reconstruct the morphology of glial cells and extract cells morphometric characteristics. Based on these characteristics, *NutriMorph* clusters the microglia into different morphological subpopulations across control (healthy mice) and high-fat diets (obese mice) feeding conditions. Its performance was tested on the microglia image dataset of Cansell et al., 2021, which results, mainly based on inflammatory factors expression levels in the hypothalamus, are concomitant with our hypothesis. We show that early hypothalamic inflammation could be already set on within a few hours through modification of microglia subpopulation proportions, instead of a couple of weeks and months. Our results tend to confirm the propensity of microglia in the ARC to branch after only a few hours of HFD diet. Furthermore, *NutriMorph* use can be broaden to the image analysis of other cells with processes that undergo morphological changes.

## 2. Material and methods

### 2.1. Algorithms and data availability

The *NutriMorph* project (written in Python v.3.8.0) is available at https://gitlab.inria.fr/edebreuv/nutrimorph under CeCILL license. To access microscopy data and the full results of their analysis with *NutriMorph*, contact the corresponding author.

### 2.2. Experimental protocols

As this paper is an extension of previous work, the protocols for animals, feeding and immunohistochemical analysis were described as in Cansell et al., 2021.

#### 2.2.1. Animals

8-week-old C57Bl/6J male mice (20–25 g, Janvier Labs, France) were group housed (two animals/cage) in a room maintained at 22 ± 1°C with a reversed 12-hour light / 12-hour dark cycle and were acclimatized for 2–3 weeks before experiments were performed. Animals had access to water and standard diets (high carbohydrates, CHO) ad libitum (3,395 kcal/kg with 25.2% from proteins, 13.5% fat, and 61.3% from carbohydrates; Safe #A03). All protocols were carried out in accordance with French standard ethical guidelines for laboratory animals and with approval of the Animal Care Committee (Nice-French Riviera, registered number 04042.02; University Paris Descartes registered number CEEA34. EA.027.11 and CEEA16-032; APAFIS#14072-2018030213588970v6).

#### 2.2.2. High-fat feeding protocols

All mice were food-deprived for 2 hours prior the onset of the dark cycle to synchronize groups. The 2-hour food deprivation at the end of the light period does not alter metabolic state as evidenced by hormonal examination (Nuzzaci et al., 2020), and maintains natural neuroendocrine control of food intake, based on synergic action of peripheral hormones (Matarazzo et al., 2012). Before food deprivation, bedding was changed to remove any leftover food in the bottom of the cage. At the beginning of the dark cycle (T = 0 hour) mice were fed either standard diet (SD, Safe #A03) or HFD (4,494.5 kcal/kg with 16.1% from proteins, 40.9% from fat, and 43% from carbohydrates; Safe #U8954P V0100). The non-fed group used for normalization was killed on the same day at the beginning of the dark cycle (T = 0 hour) and had unlimited access to SD from their arrival to the laboratory until the beginning of experiments. Then, 1, 3, and 6 hours after food exposure, food intake was measured. Animals from each of the 1-, 3-, and 6-hour groups were different.

#### 2.2.3. Immunohistochemical analysis

Mice are first i.p. injected with 100 mg.kg-1 pentobarbital sodium for terminal anesthesia. Brains were harvested from mice perfused with 4% paraformaldehyde in phosphate buffer saline (PBS) and postfixed in the same fixative overnight at 4°C. Brain coronal sections (30 μm) were cut on a vibratome, blocked for 1 hour with 3% normal goat serum in PBS containing 0.1% Triton X-100 and incubated with primary antibodies overnight at 4°C. Primary anti-rabbit antibodies were against Iba1 (1:500, CP290A, B, Biocare Medical) and GFAP (1:300, Z0334, Dako, Denmark). Adequate Alexa Fluor 488 conjugated secondary antibodies were used for immunofluorescence microscopy. Sections at −1.70 mm relative to Bregma were mounted in VECTASHIELD solution (H-1000, Vector Laboratories). 3D mosaics of 1024 × 1024 images were acquired with a TCS SP5 laser-scanning confocal microscope (Leica Microsystems, Nanterre, France) through a ×40/1.4 oil immersion objective for GFAP staining and ×63/1.4 oil immersion objective for Iba1 staining, with a z-step of 2 μm.

### 2.3. Image processing and analysis methods

#### 2.3.1. Image data

The image dataset from Cansell et al., 2021 consists in images of the hypothalamic ARC for two experimental conditions: (1) CHO for mouse fed with standard diet (high carbohydrates), (2) DIO for mouse fed with HFD. Each condition is divided into 1- hour, 3-hour and 6-hour durations of feeding. For each condition, about 10 3- dimensional images are acquired with size 1024 x 1024 x 15 (pixel x pixel x z-stack).

#### 2.3.2. Image preprocessing and analysis for soma detection

Image preprocessing for soma detection involves local intensity normalization, hysteresis thresholding (Fu and Mui, 1981), object cleaning using mathematical morphology (Haralick et al., 1987) and measurement-based outlier removal (https://scikit-image.org/). The detected cells, corresponding images and their characteristics (Table S1).

#### 2.3.3. Image preprocessing and analysis for processes detection

Due to their tubular structures, processes are first enhanced using the Frangi method ((Frangi et al., 1998) ; https://gitlab.inria.fr/edebreuv/frangi3). Histogram equalization (adaptative or not), hysteresis thresholding, object cleaning using mathematical morphology and measurement-based outlier removal, and skeletonization are then carried out (https://scikit-image.org/). The detected extensions, corresponding images and their characteristics (Table S1).

To determine the c image intensity-dependent parameter necessary for Frangi vesselness enhancement, a GUI interface was created (available in the GitLab project at NutriMorph > brick > processing > frangi3gui.py). The algorithm takes 5 arguments: (1) the path of the image, (2) the MIP axis – generally the Z axis, (3) the bright on dark option, (4) the X and (5) Y coordinate for cropping the image and allowing a faster interface experience.

#### 2.3.4. Connecting the soma to the extensions and filling gaps in processes

Because of heterogeneous staining, varying extension diameters, and noise, the detected extensions are fragmented, and they might not be connected to their corresponding soma. The shortest paths between some source and target points in a specific image intensity-depend metric is used to reconnect the fragments together, and to link the dangling extensions to their soma. We used the well-known Dijkstra shortest path algorithm ((Dijkstra, 1959) ; https://gitlab.inria.fr/edebreuv/dijkstra-img). The image intensity-depend metric is such that it costs less to cross a voxel of high intensity while the cost of crossing dark voxels is prohibitive. In a first phase, the fragments close enough to a soma are connected to it. The source points are fragment end points, and the target points are the soma contour points. Once connected, they become extensions. In a second phase, the extensions are iteratively grown by reconnecting close by fragments. The source points are still fragment end points, and the target points are the close by points of the (growing) extensions. When no such shortest path has a reasonably low cost, the reconnection procedure ends.

#### 2.3.5. Extraction of graphs from the processed images

SKLGraph module (https://src.koda.cnrs.fr/eric.debreuve/sklgraph) was used and adapted to extract graphs from the detected cells on 3D images.

### 2.4. Cells features dataset

113 features were measured based on the reconstructed microglial cells (Table S1). Five features are only soma-related and measured based on the soma volume, its convex hull and its best fitting ellipsoid (http://www.juddzone.com/ALGORITHMS/least_squares_3D_ellipsoid.html). The other features are measured based on the graphs representing processes, such as the number of edges and nodes. Branches length, thickness, volume, curvature and torsion are transcribed in the table through their minimum, maximum, mean, median, standard deviation and entropy. Length and curvature distribution histograms are also computed. Branches are separated into two kind of extensions: primary (P), linked to the soma, and all the others, secondary (S) . The features are stored under CSV format and contain 558 detected cells.

### 2.5. Statistical analyses and machine learning methods

Statistical analyses of the features’ dataset were performed using pandas Python module (https://pandas.pydata.org/). Most relevant features were selected based on three criteria: (i) their pairwise correlation coefficient, the p-value of the (ii) Kolmogorov-Smirnov and (iii) Wilcoxon signed-rank tests between CHO and DIO conditions. Principal Component Analyses (PCA) were performed on the selected features, and dimension reductions to 2 or 3 dimensions were studied. k-Means algorithm was carried out with k equal to 2, 3, and 4, to detect potential proportion shifts between plausible cells subpopulations. Histograms were computed for each experimental duration showing the proportion of each k-means label per experimental condition (CHO and DIO). Bhattacharyya distance (Bhattacharyya, 1946) was calculated between the histograms for a given duration. As an alternative to histograms, kernel density estimators depending on labels, conditions and/or duration were also studied.

## 3. Results

### 3.1. NutriMorph

*NutriMorph* is a fully automatic neural cells tracing and analysis method for fluorescent microscopy images. It reconstructs the cells soma and processes and performs a statistical analysis and clustering of these cells. Algorithm 1 (Figure 1A) depicts the main algorithmic steps of the method. *NutriMorph* can either be fed with a single image or a directory of images. In the first case, no statistical analysis or clustering is performed and the output only comprises a CSV file with each detected cell’s features. For directory input, *NutriMorph* runs an image analysis function for each image, and once done, returns a global CSV file containing all the features for each detected cell in each image, labeled by their specific experimental condition. In detail, the image analysis pipeline (Algorithm 2, Figure 1B) detects the soma of the cells, their extensions, while dealing with reconnecting somas and extensions, and reconnecting extension fragments together if needed. Once the cell objects are complete, their representation as a graph is computed to turn their topological information and 3-dimensional (3D) geometry in a form adapted to computational analysis. Selected topological and geometrical characteristics of the cells are measured on these graphs and stored in a features file in CSV format.

**Figure 1.**
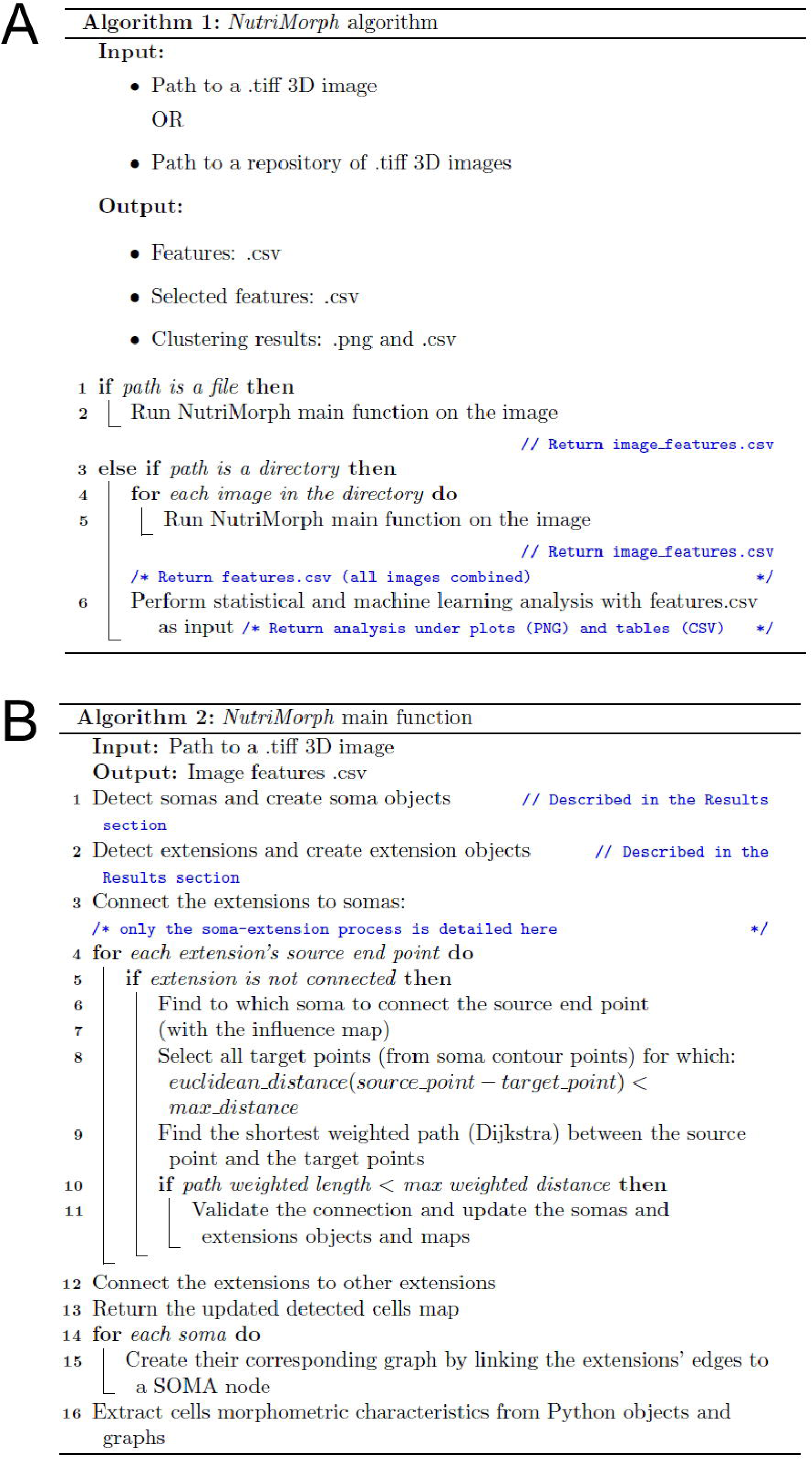
Algorithm 1, *NutriMorph* and Algorithm 2, *NutriMorph* main function1. The NutriMorph tool has been implemented in Python 3 and is available at: https://gitlab.inria.fr/edebreuv/nutrimorph

The behavior of *NutriMorph* is guided by some biological parameters, that are specified in micron units for convenience, such as the minimal and maximal size (volume) of a soma, or the maximal length of the processes. The algorithm takes into account that the 3D images may have, and usually do have, different resolutions in X, Y and Z directions when converting quantities from pixels/voxels into microns and vice versa.

#### 3.1.1. Soma detection

The input 3D image is first normalized on a min-max scale between 0 and 1:

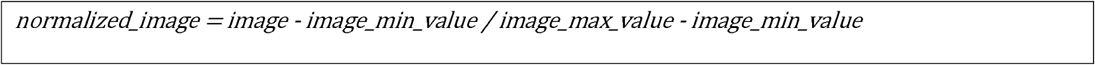

as to allow for a consistent contrast despite images being potentially acquired in various experimental conditions. The normalized image is then preprocessed in order to denoise it and detect somas structures. Hysteresis thresholding (low = 0.1500, high = 0.7126) is performed. Opening and closing (structural element radius = 0.481 microns) are conducted on the resulting image to remove small, spurious elements, and to fill small holes inside the somas and smooth their somas contours, respectively. The volumes of the somas are calculated based on their number of voxels. Somas with an excessively small (< 116 micron3) or big volume (> 1160 micron3) – specified in the *NutriMorph* configuration file – are deleted from the image. Note that this step complements the opening operation above since opening does not account for the total volume. The somas are then labeled, and a 3D co-called influence map is created (Figure 2A). This map stores at each voxel location the label of the closest soma. The soma objects are enriched with characteristics such as their volume in micron, their centroid and the position of their contour points.

**Figure 2.**
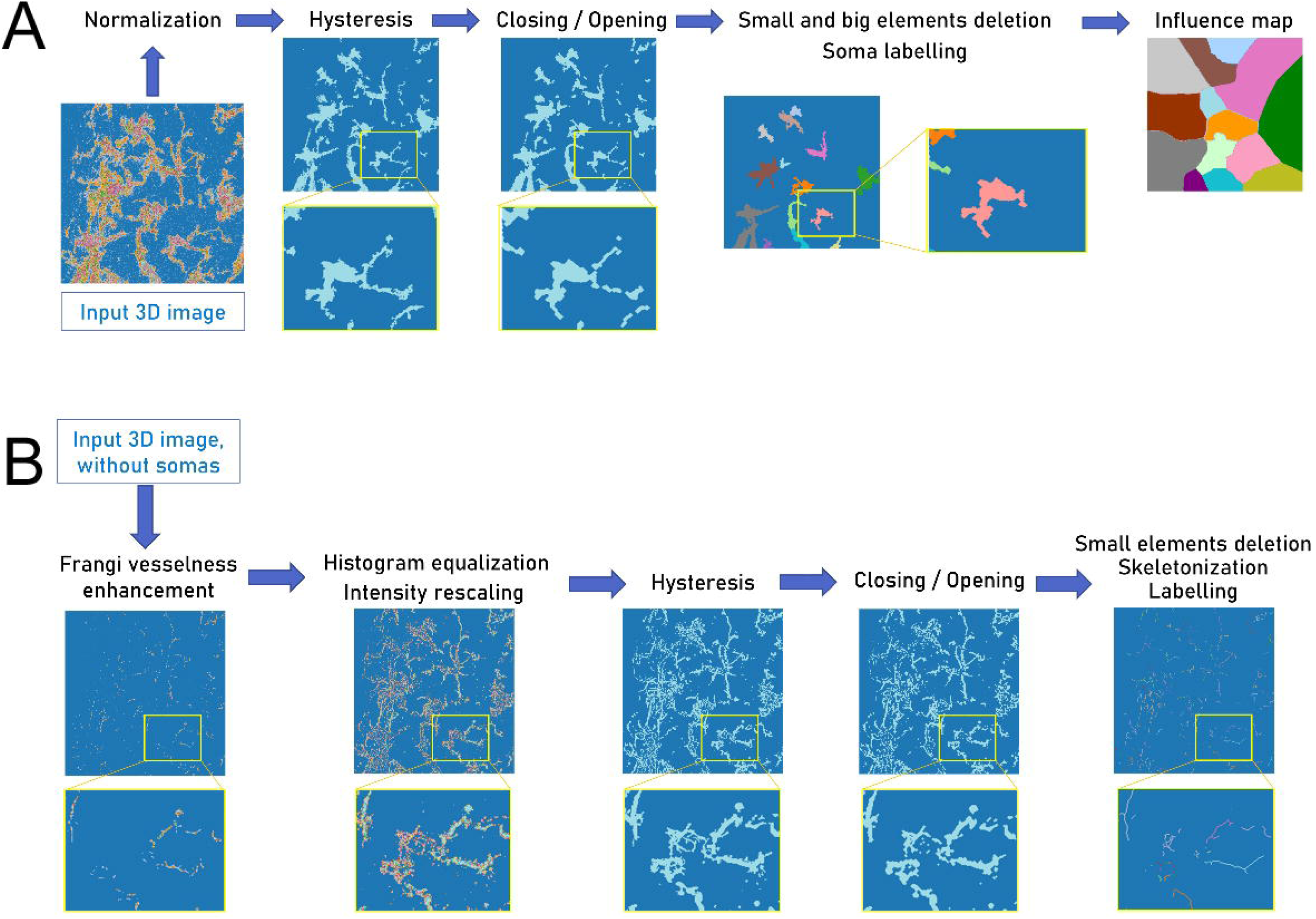
Soma and processes detection. A. Soma detection. All the 3D images are represented in 2D after maximum intensity projection (MIP) in the Z axis. The background is represented in azure-blue. The input 3D microscopy image is first normalized between 0 and 1. Hysteresis thresholding and closing/opening are performed on the image to highlight cellular structures, binarize the image, and remove noise. Small and spurious elements as well as voluminous merged cells are removed, and somas are labeled. A soma influence map is then created. B. Processes detection. All the 3D images are represented in 2D after maximum intensity projection in the Z axis. The background is represented in azure-blue. Voxels corresponding to somas are deleted from the input 3D microscopy image. The Frangi method is performed to enhance the tubular structures in the image. To better enhance all the processes, histogram equalization and intensity rescaling are carried out. Processes are binarized by hysteresis thresholding and cleaned by closing/opening. Small elements are then removed. The processes are skeletonized and labeled.

#### 3.1.2. Processes detection

Detected fragments of processes are called extensions. First, voxels corresponding to somas are set to 0, to avoid detecting erroneous extensions. Tubular structures are then enhanced using the Frangi vesselness method in 3D (Frangi et al., 1998). The following parameters were selected: scale range = (0.1, 3.1), scale step = 1, alpha = 0.8, beta = 0.5, bright on dark variant. To determine the c parameter, which is dependent on the image intensity distribution, an interactive Graphical User Interface (GUI) was created (see Methods). The Frangi method relies on the eigenvalues of the image second order derivatives (Hessian matrix) to compute the likeliness of a region to contain vessels, or more generally, tubular structures. The second order derivatives were computed using finite difference approximations. The Frangi3 filtering produces an image where the tubular structures are emphasized while other structures are dimmed. It also returns a scale map that represents the approximate thickness of the tubular structures, i.e. the extensions in our case. These structures are further enhanced with histogram equalization. The enhanced image is than processed through hysteresis thresholding (low = 10 x minimum_non-zero_intensity, high = 0.6 x maximum_intensity) to detect the (possibly fragmented) extensions.

Morphological closing and opening (circular structural element with radius = 0.24 microns) are then performed. Small extensions (< 5.7 microns in length) are removed. The remaining extensions are then skeletonized (Lee et al., 1994) and labeled (Figure 2B). The extension objects are enriched with characteristics such as their thicknesses along the extension (Frangi scale), their length in micron, and their end points.

#### 3.1.3. Connection of extensions to soma and reconnection of extension fragments

In microscopy images, glial processes are heterogeneously marked by the fluorescent agent in microscopy images, they vary in diameter, and their appearance is degraded by noise. In consequence, their detection usually produces several elongated fragments for one process. To reconnect the fragments together to recover the processes, and the processes to their somas to recover whole cells, we perform an image-driven gap-filling using a shortest-path discovering-method known as Dijkstra (Dijkstra, 1959). A connection between two fragments or between a fragment and a soma is defined as the path of minimal length in a modified, image-based metric joining a fragment end point to either a contour point of a soma or a point of another fragment (not necessarily an end point). To limit the search space, only close by points are considered for connection.

This connection procedure is divided into two steps (Figure 3): (i) the connection between somas and nearby extensions or extension fragments not yet connected to a soma, and (ii) once done, the connection between extensions already attached to a soma and nearby fragments to reconstruct the whole processes. Step (i) is iterated until no extension or fragment can be connected to a soma by a path whose length is below a selected threshold. Step (ii) is iterated until no fragment can be connected to an extension by a path whose length is below a selected threshold. The modified, image-based metric used to compute path lengths is defined by a so-called cost map giving the cost of passing through each voxel. For the voxels of the somas and the extensions, or the extension fragments, this cost is set to (the computer representation of) infinity to prevent any path from crossing an object. To favor paths across bright areas, which are more likely to correspond to actual extension pieces, the cost of the other voxels is set to the inverse of their intensity (with a safeguard to avoid divisions by zero):

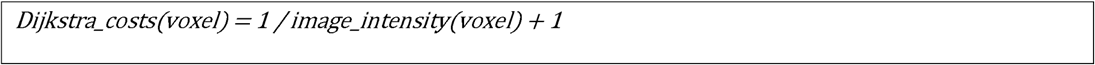

**Figure 3.**
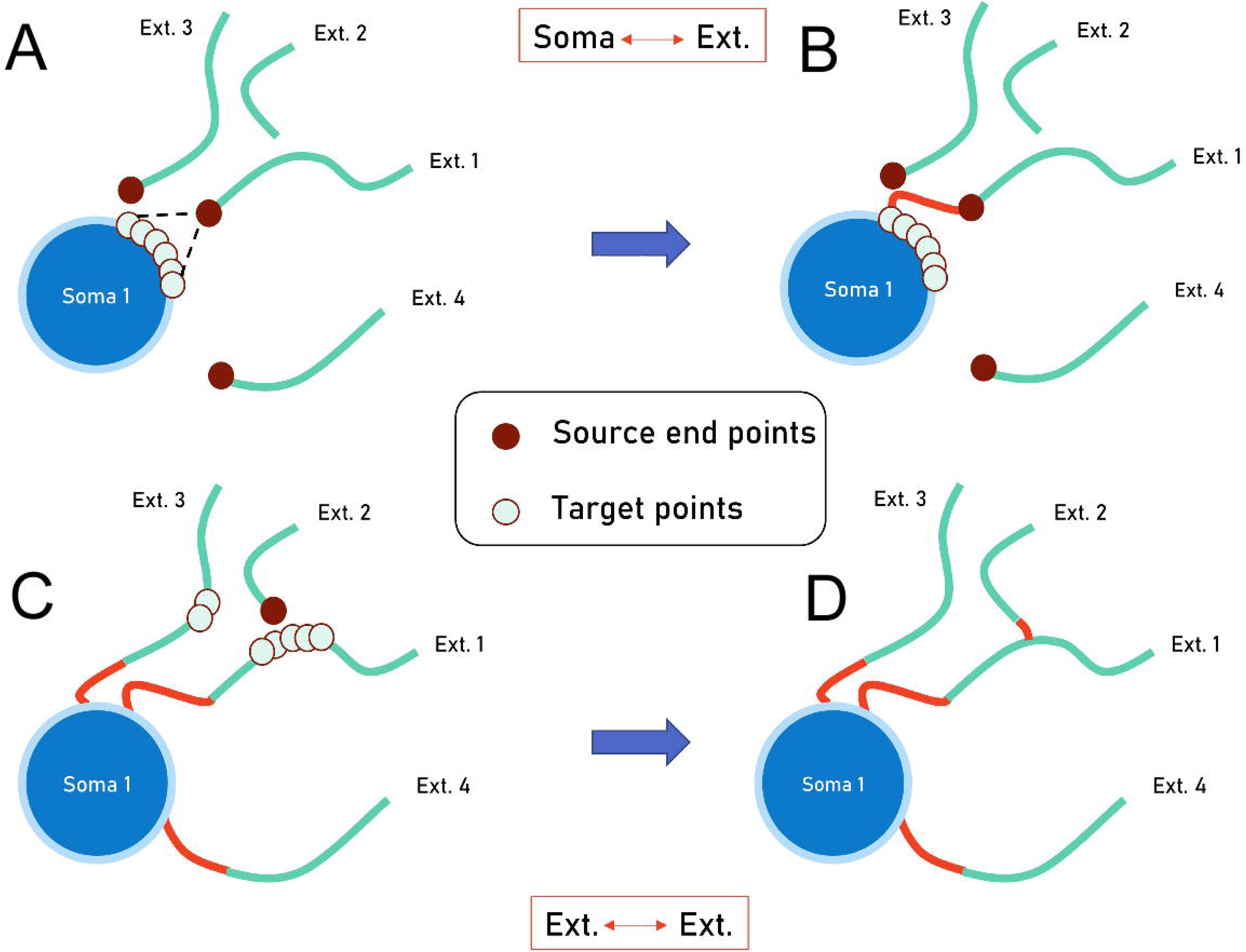
Connection between somas and extensions, and in-between extensions. Somas-Extensions connections. A. For each soma, source end points for connection are found in the extensions within the soma influence. For each source point, all the potential target points are found in the contour points of the target soma such as their Euclidean distance is smaller than a defined maximum distance. B. A shortest weighted path is then searched between the source points and all the target points together with the Dijkstra algorithm. If the weighted path length is inferior to a certain maximum weighted length, then the connection is validated and soma objects, extension objects and maps are updated. C. Extensions-Extensions connections. D. The same process is performed, but target points are on extensions, and may not be restricted to a single extension.

After each new connection, the voxels of the connection path are set to “infinity” in the cost map since they now form part of an extension. Note that the immediate neighborhood of the somas and/or the extensions and the fragments are also set to infinity in the cost map (beside exceptions) to avoid the connection paths passing to close (being tangent in particular) to existing structures. As exceptions, the immediate neighborhood of the soma with its extensions and the extension or fragment considered for connection are not set to infinity, otherwise no connections could be made.

As mentioned above, only close by points are considered for connection. It allows to reduce the computational load drastically. But it also encourages likely connections. This closeness is defined by a threshold on the Euclidean distance between candidate end points for a path (set to 217 microns) where the candidate points must both belong to the same region of a so-called soma influence map. This map is computed as the Voronoi diagram (Aurenhammer, 1991) of the somas with their extensions, meaning that a region of it describes the set of points closer to a given soma+extensions than to any other soma+extensions. After each new connection (whether extension or fragment to soma, or fragment to extension), the influence map is recomputed to account for the new configuration.

This whole procedure produces a set of glial cells (somas with their complete extensions) in the form of a labeled map where each cell has its own label. The extension fragments that were not connected to a soma or an extension are considered as spurious objects and discarded.

#### 3.1.4. Extraction of glial cells graphs from the 3D labeled maps

Once all the cells are detected in the 3D image, they are converted to graphs, i.e. a set of nodes connected by edges (Figure 4). There are four types of nodes: nodes that represent somas’ center of mass, nodes that correspond to the anchor points between a soma and its extensions (they are root nodes and are connected to the center of mass node), nodes that correspond to an extension branching point where it splits into two “daughter” extensions, and nodes that correspond to extension end points. Each edge stores details such as the list of voxels forming the corresponding extension piece, the thicknesses, curvature, etc along the piece, … In the following study, primary extensions are defined as the edges between a root node and the first branching point of an extension (if any; otherwise, the whole extension). The remaining edges are called secondary extensions.

**Figure 4.**
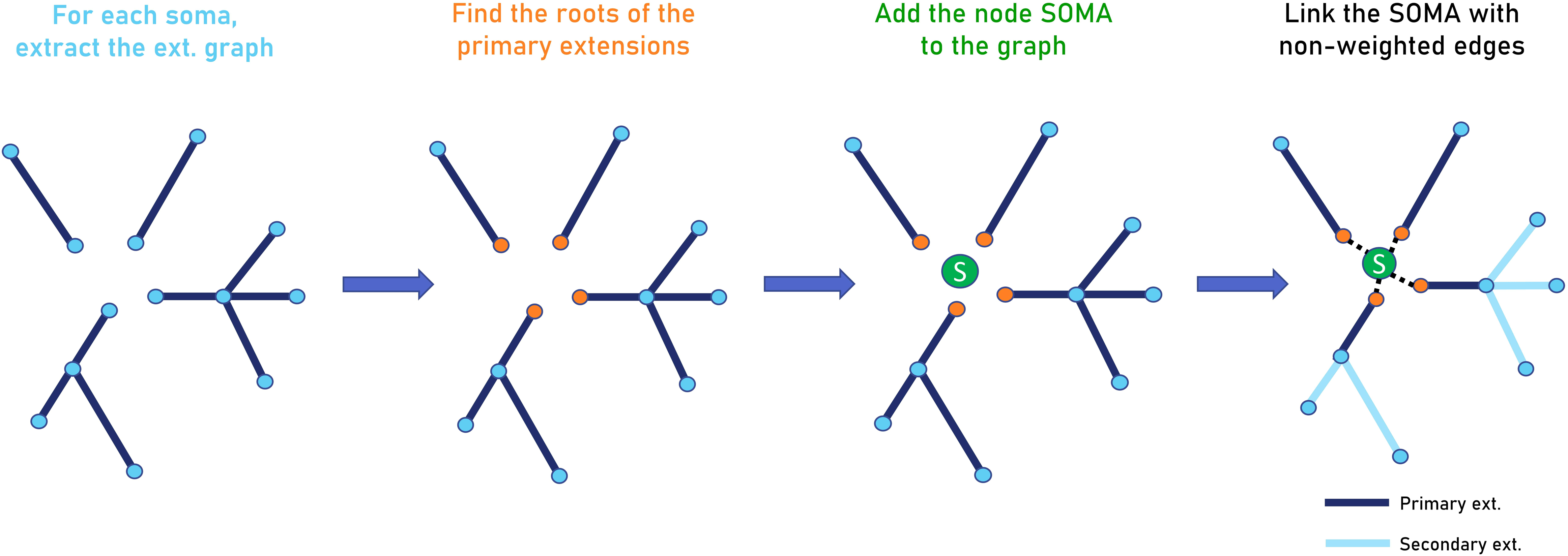
Pipeline of the conversion of detected glial cells into graphs. First, somas are removed from the labeled cells image while keeping track of their associated processes. The processes are converted to graphs after skeletonization by SKLGraph. Then, the roots of the primary extensions are found. The processes are then linked to a special SOMA node, positioned at the soma’s centroid, with unweighted edges to their root point. Primary extensions are defined as the ones linked to the soma through a root node, and the secondary extensions are all the others.

#### 3.1.5. Calculations of cells’ morphometric features

Several features are calculated to describe the soma morphology. The convexity of the soma is described by the ratio between the soma volume and the volume of its (3D) convex hull. The overall shape (e.g., whether it is rather flattened in a certain direction, or more spherical) is expressed by the ratios between the semi-axes of the best fitting ellipsoid, i.e. 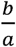 and 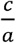, if a, b, and c, with a > b > C, are the semi-axes. The orientation of the cells in the image is measured by the two couples of spherical angles (ϕ, θ) for the axes corresponding to *a* and *b*.

The processes features are the minimum, maximum, mean, median, standard deviation and entropy of the following edge characteristics: length, thickness, volume, curvature and torsion. These features are computed in three contexts: for the primary extensions, for the secondary extensions, and for the extensions as a whole. In addition, the number of branching nodes, the number of total, primary and secondary edges, the total length of the extensions (per soma) and the histograms of edge length and curvature are added to the features.

These features, along with image names, soma labels and experimental conditions and durations, were stored in CSV format (Table S1).

#### 3.1.6. Statistical analysis and cell clustering

In this study of obesity, two conditions were represented: CHO (Chow) for mice fed with standard diets for given durations (1 hour, 3 hours, 6 hours) and DIO (Diet- Induced Obesity) for mice fed with high-fat diets for the same durations. Only scalar features (i.e. letting aside histograms and spherical angles) were integrated in the following statistical and machine learning procedures.

The most relevant cells features are selected based on three criteria: (i) the correlation coefficient between feature pairs (mixed conditions; represented on a heatmap, Figure S1), the p-value of the (ii) Kolmogorov-Smirnov two-sample and (iii) Wilcoxon signed-rank two-sample tests between CHO and DIO conditions. The Kolmogorov-Smirnov two-sample test permits to compare feature distributions between each condition, while the Wilcoxon signed-rank two-sample test does a median comparison. Features with pairwise correlation coefficient higher than 0.9, p- values (between CHO and DIO, all durations together) higher than 10^-2^ and p-values (between CHO and DIO at 6 hours) higher than 10^-3^ were removed.

A Principal Component Analyses (PCA) was additionally performed on the selected features. However, no dimension reduction appeared suitable for our purpose.

k-Means clustering was then carried out with k equal to 2, 3, and 4, to discover if distinct subpopulations exist, whether within or between the two conditions, and if yes, if their proportions changes between the conditions. Note that the number of sub-populations might or might not be equal to the number of conditions (CHO and DIO). We studied histograms of the cluster sizes for each value of k, each experimental duration, and each experimental condition (CHO and DIO). The Bhattacharyya distance (Equation S1) was calculated between the CHO and DIO histograms for a given duration to determine their potential evolution as a function of time. Features distributions depending on clusters, conditions and/or duration can also be drawn.

Note that density-based or model-based alternative clustering methods, to name a few, such as DBSCAN (Ester et al., 1996) or Gaussian Mixture Model (Xu and Jordan, 1996) exist. However, our number of samples is not large enough for such methods.

### 3.2. Results and performance evaluation

*NutriMorph* was tested on a set of fluorescent microscopy images of microglia of the hypothalamic arcuate nucleus, a brain region particularly affected by obesity-induced neuroinflammation, and which regulates feeding behavior, including satiety. The images were acquired for two different conditions: mice fed for 1 hour, 3 hours, and 6 hours with a standard diet (CHO) and mice fed for the same durations with a high-fat diet (DIO). The mice were different for each duration and condition.

The goal of the study is to highlight a possible microglial activation within the first postprandial hours, way before the excess bodyweight onset.

#### 3.2.1. Microglia get transiently activated in two different morphological subpopulations in the arcuate nucleus of the hypothalamus after a few hours of HFD feeding

Various characteristics were extracted from all the detected cells in all images with *NutriMorph* (Table S1). After features selection based on statistical tests (high-correlated features dropping, Kolmogorov Smirnov two-sample test and Wilcoxon signed-rank two-sample test, see Materials and Methods), 2D and 3D PCAs were implemented (Figure S2). No clear separation of the data was visible, even for the 6- hour CHO and DIO groups. For our dataset, the PCA-based dimension reduction was not appropriate, with an explained variance by the first principal components (PC) being pretty low (explained variance for PC1 = 26%, PC2 = 11%, PC3 = 9%). Additionally, in the reduced 2D or 3D dimensional spaces, there was no visible distinction between the two conditions, which confirms the inappropriateness of a PCA. Interestingly, this observation might be explained by the presence of multiple morphological subpopulations of microglial cells across conditions and/or durations.

To detect such potential subpopulations within conditions, k-means clusterings were carried out with a number of clusters ranging from 2 (Figure 5) to 4 (Figure S3). The per-cluster proportions in the CHO and DIO conditions for each condition and duration are presented as barplots in Figure 5. On per-condition barplots (all durations together), no clear difference in proportions is observed. For instance, for k = 2, there are 35% of cluster 0 for CHO and 39% for DIO (Figure 5A). This confirms our conclusion regarding the lack of interest of PCA here. However, from a temporal perspective, still for k = 2 (Figure 5B), a notable difference is observed for the DIO condition between the cluster proportions between 1 hour and 3 hours: cluster 0 changes from 36% to 24% and cluster 1 changes from 64% to 76%. Then, from 3 hours to 6 hours, an inversion of the proportions occurs: cluster 0 changes from 24% to 54% and cluster 1 changes from 76% to 46%. In the CHO condition, the clusters exhibit similar proportions throughout time. The results for k equal to 3 and 4 follow similar trends, with some clusters being almost nonexistent (Figure S3). These distinctive temporal evolutions in the proportions of the clusters, i.e. cell subpopulations, can also be characterized by the Bhattacharyya distances (Equation S1) between the CHO and DIO conditions at each duration (Figure 5C).

**Figure 5.**
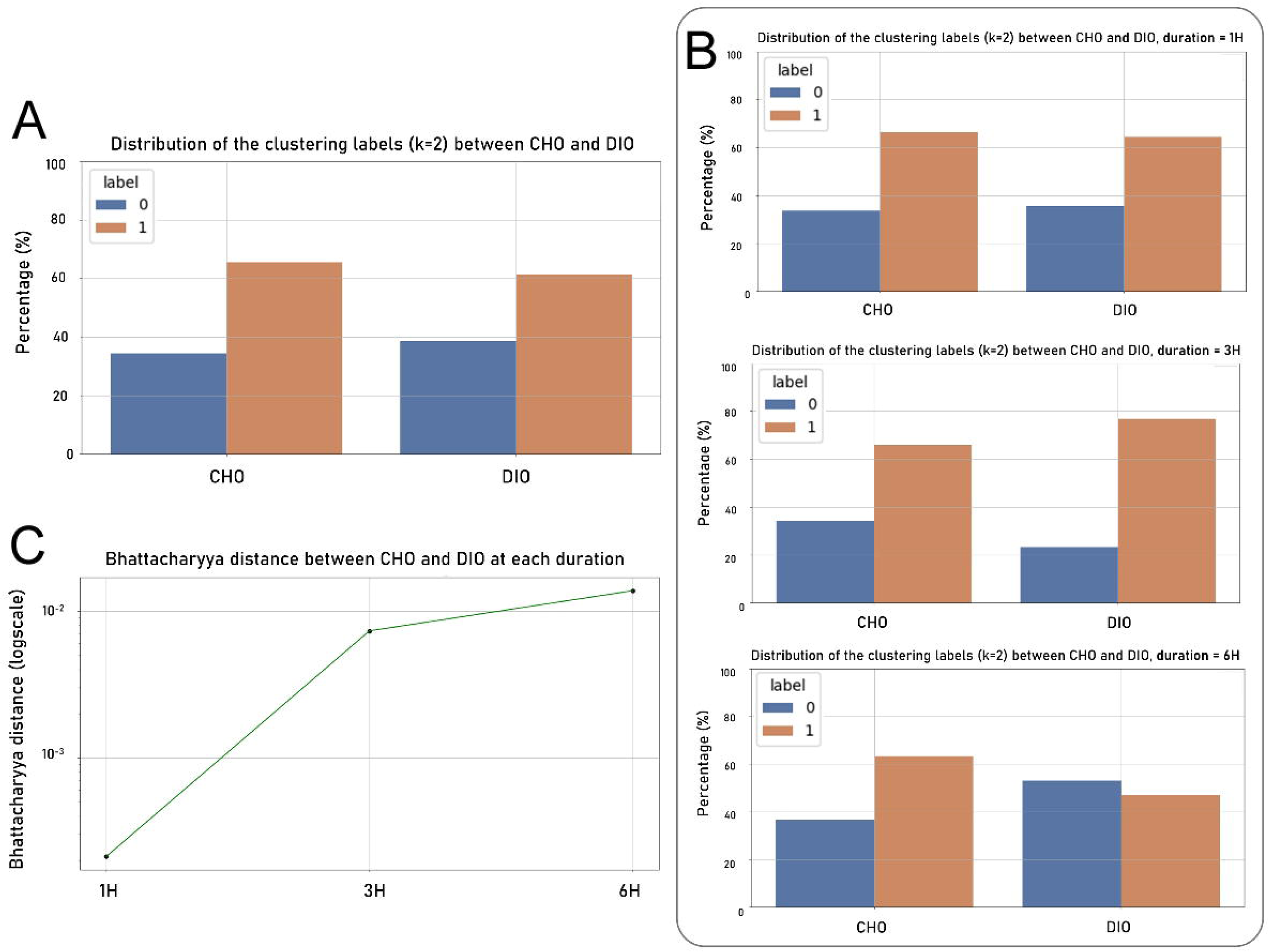
k-Means (k=2) clustering results for each diet (CHO and DIO). A. Distribution of the clustering labels between CHO and DIO conditions. The percentage of cells labeled 0 or 1 for each condition is represented. When durations are not differentiated, little difference is observed in the proportion of the labels 0 and 1 between CHO and DIO. B. Distribution of the clustering labels between CHO and DIO for each experimental duration. When durations are differentiated, the proportions only change in the case of the DIO condition. Between 1H and 3H the proportion of cells labeled 1 is increasing (from 64% to 76%), and a shift in proportion is then observed between 3H and 6H (from 76% to 46%). C. Bhattacharyya distance between the conditions CHO and DIO at each duration. The difference between histograms of CHO and DIO conditions was computed for each duration. The difference between CHO and DIO increases in the first postprandial hours.

In light of this analysis, we hypothesize that two main morphological subpopulations of microglia are present and that, in response to HFD short-term exposition, transient modification of the microglial morphology occurs. This is concomitant with published results of transient liberation of inflammatory mediators by these cells after a few hours HFD feeding (Cansell et al., 2021).

#### 3.2.2. Transient activation of microglia is characterized by specific morphological markers

We plotted the distributions of the features (computed using the kernel density estimator; KDE) for each cluster (k=2), all durations together but separately for the CHO and DIO conditions (Figure 6). The distributions appear different between the two clusters, with two situations: their variance is similar in the two clusters, but their average is different (Mean_length_P and Mean_volume_P), and both the variances and the averages are different (most notably Total_ext_length_P). The distributions of the less easily interpretable features can be found in the Supplementary Material. Some of them also appear different between the two clusters, particularly the ones based on entropy. These differences in distribution allow us to build a typical sketch for each cluster, or subpopulation. The cells in cluster 1 have approximately 4 times more extensions, which decomposes into a 1.5-fold for primary extensions and a 6- fold for secondary extensions compared to cluster 0. The mean curvature of their extensions, especially the primary ones, is higher. The length of their extensions is about 1.5 micron higher for primary extensions and 1 micron higher for secondary extensions. The mean volume of their extensions is also about 0.3 µm^3^ higher for primary extensions and 0.1 µm^3^ higher for secondary extensions).

**Figure 6.**
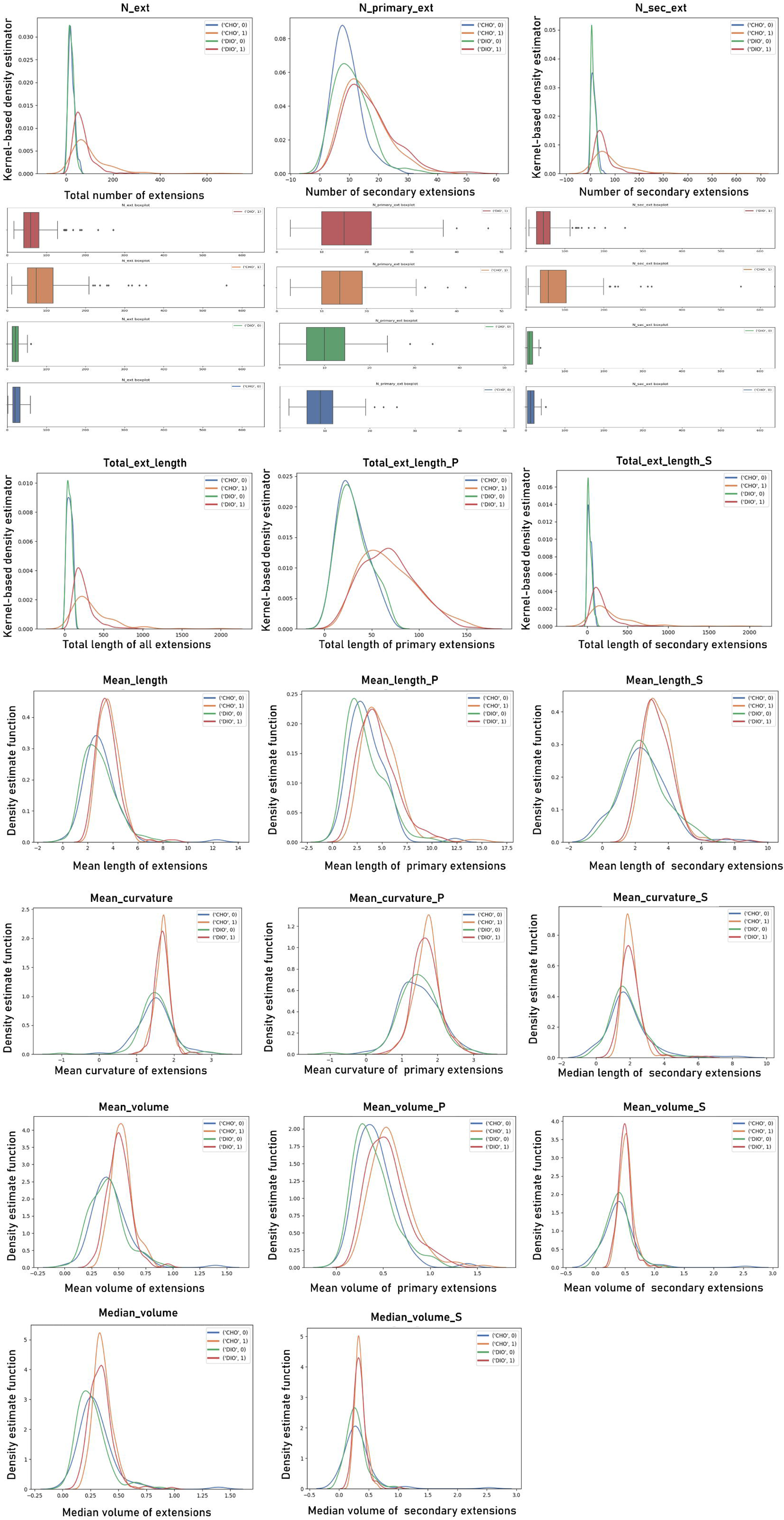
Kernel-based density estimators for different features, differentiated by their condition (CHO, DIO) and their k-Means clustering label (0, 1). Kernel-based density of each feature was computed based on each feature’s histogram. Boxplots for the three first features are shown. The density estimators show clear separation of the two clustering labels and similar distributions for the conditions with the same label. This suggest relevant morphological difference in between the two identified clusters, a.k.a. microglial subpopulations. For explanation of the features’ name, see Table S1. Boxplots for the presented estimators are represented in Figure S4. The density estimator and the boxplots of the other features can be found in the Supplementary Materials folder. Some of these other features, although less comprehensive, show significant difference between the density estimator for label 0 and 1.

Based on biological knowledge, we thus hypothesize that cluster 1 corresponds to a subpopulation of activated microglia, while cluster 0 represents a quiescent subpopulation. Therefore, in DIO, the increase of quiescent microglia proportion at 3 hours may be due to a recruiting of glial cells and/or an endocrine state, while the inversion in proportion at 6 hours, would denote a greater number of activated microglia in the ARC, that will influence neuronal functioning.

## 4. Discussion and conclusion

We presented *NutriMorph*, an automatic method of detection and analysis of glial cells, in fluorescent microscopy images. It comprises the detection of the somas and their multibranched processes, the conversion into a graph representation and the computation of an extensive number of morphological characteristics. Through statistical analysis and machine learning, *NutriMorph* allowed to compare different experimental conditions in terms of cell morphology.

Once the parameters of the method have been chosen, *NutriMorph* can process and analyze a set of images in batch, i.e. without user intervention.

Further improvements of the methodology may include an automatic selection of the optimal Frangi parameter c, alternative statistical and machine learning methods in light of other results on medium-term and long-term experiments (see below), and an inclusion of non-scalar features in the clustering step.

Regarding our biological objective, we showed that microglia get transiently activated after only a few hours of high-fat diet feeding, which translates into two different morphological subpopulations in the arcuate nucleus of the hypothalamus. One subpopulation is identified as quiescent-state microglia while the other one is activated, where activation is characterized by longer, larger and more branched processes. Very short-time proportion changes are observed in the two subpopulations in case of obesity. At 3 hours after HFD feeding, less microglia are in an activated state, which could be explained by a recruiting process or/and an endocrine state of the microglia corresponding to a quiescent-like morphology. At 6 hours after HFD-feeding, the proportions inverse and more microglia are in the activated state, and they might interact more intensively with the hypothalamic neuronal component that regulate feeding behavior.

This early activation, although supposedly reversible, let us wonder whether regular HDF intake in youth could dramatically foster obesity and its comorbidities in growing adults, due to neural plasticity. To have a better insight on this problematic, further analysis could be performed usint *NutriMorph* on data for longer HFD-exposition durations (days, weeks and months). For these medium-term and long-term experiments, we expect to see less transitory shifts in proportions of the subpopulations, but greater differences between the standard diet condition and the high-fat diet one. This analysis could be conducted in two modes, namely with or without taking into account the spatial distribution of the sub-populations between the arcuate nucleus and the median eminence, another very affected region. These results would pave the way for pharmacogenetic investigations to determine whether inhibition of early postprandial activation of glial cells could reduce food intake and prevent obesity to be able to offer innovative therapeutic management for the treatment of obesity.

## Supporting information

Graphical Abstract

## Supplementary figures and tables caption

Figure S1. Pairwise correlation heatmap of the cells’ features. The redder the intersection points in between two features, the more correlated they are.

Figure S2. 2- and 3-dimensional Principal Components Analyses (PCA). A. 2D PCA differencing the experimental durations (1 hour, 3 hours, 6 hours) and the conditions (CHO, blue; DIO, red). B. 2D PCA combining all the durations (all data), represented in different colors, and differencing the conditions (CHO, blue shades; DIO, red shades). The two main principal components explain 26% and 11% of the variance. C. 3D PCA only differentiating CHO and DIO, for all durations. The third principal component only accounts for an additional 9% of the explained variance. Thus, dimension reduction cannot be performed on our data using PCA.

Figure S3. k-Means (A, B, C. k=3 and D, E, F. k=4) clustering results for each diet (CHO and DIO). A,D. Distribution of the clustering labels between CHO and DIO conditions. B,E. Distribution of the clustering labels between CHO and DIO for each experimental duration. C,F. Bhattacharyya distance between the conditions CHO and DIO at each duration.

These graphs closely follow the trends detailed in Figure 5, except for two points: (i) there are other cluster labels, but they are present in very low proportion, close to null; (ii) in B.3., the distance between CHO and DIO histograms at 6H is less important than the one at 1H and 3H, which is probably due to the influence of the other two labels.

Figure S4. Boxplots representing the distribution of the features in Figure 6, for each diet (CHO, DIO) and clustering label (0,1). For details on the features, see Table S1. To access all the features’ kernel-based density estimator and the corresponding boxplots, consult the corresponding folders in Supplementary Materials.

*Equation S1* Bhattacharyya distance is computed by

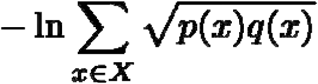

where p(x) and q(x) are two different distributions with values x belonging to the X space.

Table S1. Features extracted from the graph representations of detected glial cells.

## Author contributions

Project conception, Eric Debreuve and Carole Rovère; Biological experiments and data acquisition, Céline Cansell and Clara Sanchez; Conceptualization – algorithm, Eric Debreuve; Methodology – algorithm, Morgane Nadal and Eric Debreuve; Proof- of-concept creation – algorithm, Sarah Laroui, Eric Debreuve and Xavier Descombes; Investigation – algorithm, Morgane Nadal and Eric Debreuve; Results interpretation, Morgane Nadal, Clara Sanchez, Carole Rovère and Eric Debreuve; Writing – Original, Morgane Nadal, Clara Sanchez, Eric Debreuve and Carole Rovère; Funding Acquisition, Eric Debreuve and Carole Rovère; Resources, Eric Debreuve and Carole Rovère; Supervision, Eric Debreuve and Carole Rovère.

## Acknowledgements

We thank the microscopy facility from the IPMC part of the “Microscopie Imagerie Côte d’Azur” GIS IBiSA labeled platform, especially Frédéric Brau.

## Terminology

In biology, the cell body of a neural cell is called soma and its cytoplasmic extension are called processes.

In the NutriMorph algorithm, during the processes’ detection, detected fragments of processes are named extensions. During the conversion of detected glial cells into graph, each graph edge is either called edge or extension. Primary extensions are the extensions directly linked to the soma, whilst secondary extensions are all the others.

## Data statement

The authors confirm that the data supporting the findings of this study are available within the article and its supplementary material.

## Abbreviations

ARC: Arcuate nucleus;
CHO: Chow: High carbohydrate diets
CNS: Central neural system;
CSV: Comma-separated values
DBSCAN: density-based spatial clustering of applications with noise
DIO: Diet-induced obesity;
FA: Fatty acid;
GUI: Graphical User Interface
HFD: High-fat diets;
LoG: Laplacian of Gaussians;
DoG: Difference of Gaussians;
MSER: Maximally stable extremal region;
PBS: Phosphate buffer saline;
PSF: Point Spread Function;
SD: Standard diets;
SFA: Saturated fatty acid;
SIFT: scale-invariant feature transform;
WHO: World Health Organization.

