## Supplementary figures and images for "Computational detection, characterization, and clustering of microglial cells in a mouse model of fat-induced postprandial hypothalamic inflammation"

### Graphical Abstract

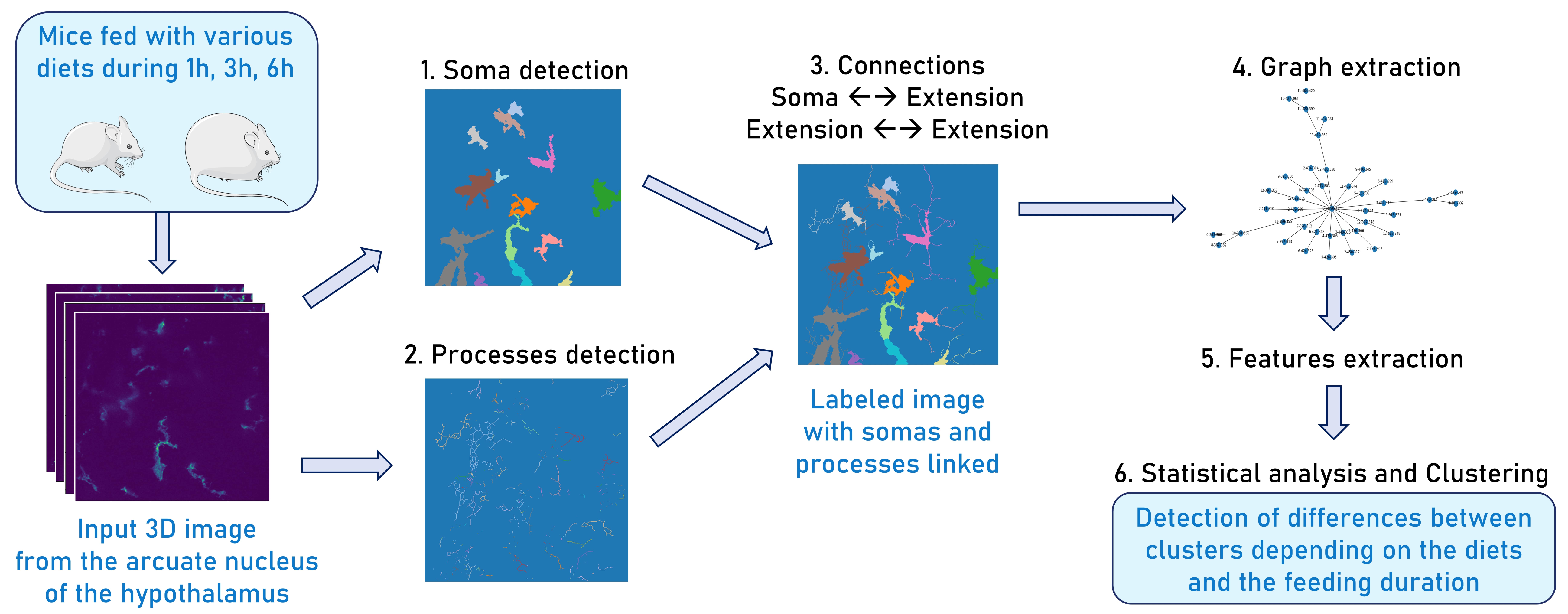

### Supplemental Figure 1

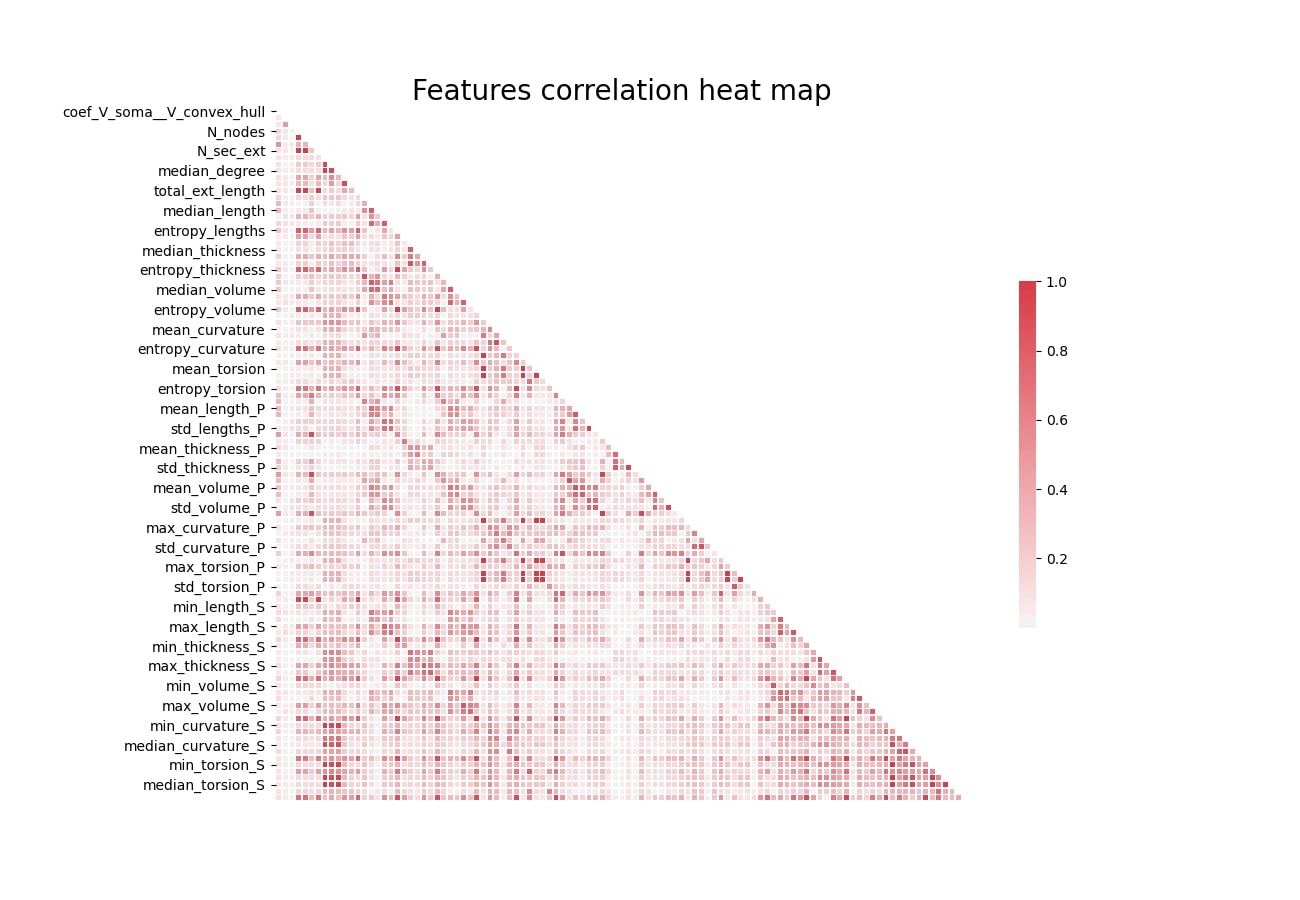
